# NLRP1B allele 2 does not respond to Val-boro-Pro (VbP) in the intestinal epithelium

**DOI:** 10.1101/2024.07.18.604161

**Authors:** Ryan J. Mazzone, Nathaniel J. Winsor, Lu Yi Li, Kristian T. Barry, Adrienne Ranger, Shawn Goyal, Justin J. Meade, Jessica Bruce, Dana J. Philpott, Jeremy Mogridge, Stephen E. Girardin

## Abstract

The intestinal mucosa must balance tolerance to commensal microbes and luminal antigens with rapid detection of enteric pathogens in order to maintain homeostasis. This balance is facilitated through the regulation of epithelial layer integrity by innate immune receptors. Certain NOD-like receptors (NLRs) expressed in intestinal epithelial cells, including NLRC4 and NLRP9B, form inflammasomes that protect against pathogens by activating caspase-1 to cause extrusion of infected cells. NLRP1B is a murine NLR encoded by five alleles of a highly polymorphic gene homologous to human NLRP1. NLRP1B forms inflammasomes in response to a variety of pathogens that cause intestinal infections, but it has almost exclusively been studied in immune cells and has not been characterized in cells of the intestinal epithelium. Here, we show that *Nlrp1b* is expressed in ileal and colonic organoids derived for C57BL/6J mice. *Nlrp1b* was upregulated by interleukin-13 in organoids and by the protozoan *Tritrichomonas muris* in vivo, suggesting that NLRP1B may be involved in defense against enteric parasites. Surprisingly, while Val-boro-Pro (VbP) activated NLRP1B in bone marrow-derived macrophages, it did not activate NLRP1B in organoids. We furthermore did not detect *Nlrp1b* in organoids derived from Balb/cJ mice, which express a different allele than the one expressed in C57BL/6J mice. Together, our results suggest that NLRP1B may have an allele-dependent function in murine IECs whose regulation is distinct from that of macrophages.

## INTRODUCTION

NOD-like receptors (NLRs) are pattern recognition receptors that detect microbe-associated molecular patterns (MAMPs) and danger-associated molecular patterns (DAMPs) in the cytosol of immune and non-immune cells, including epithelial cells ^1–3^. Upon ligand detection, some NLRs oligomerize, and recruit the adaptor protein apoptosis-associated speck-like protein (ASC), and pro-caspase-1 to form signalling complexes known as inflammasomes ^4^. This interaction typically results in the autoproteolytic processing and activation of caspase-1, which can cleave gasdermin D (GSDMD), as well as pro-inflammatory cytokines pro-IL-18 and pro-IL-1β. The liberated N-terminus of GSDMD mediates pore formation at the plasma membrane, causing inflammatory pyroptotic cell death and release of mature IL-18 and IL-1β into the extracellular space ^4^.

NLRP1 was originally characterized in human monocytes and was the first NLR known to nucleate an inflammasome ^5^. NLRP1 possesses a unique structure and is composed of an N-terminal pyrin domain followed by a NACHT domain, a leucine rich repeat (LRR) domain, a function-to-find domain (FIIND), and a caspase activation and recruitment domain (CARD) domain ^6^. NLRP1 and its murine orthologue NLRP1B, both undergo post-translational autoproteolysis at the ZU5 and UPA interface of the FIIND domain and remain non-covalently associated in an inactive state ^6^. The inactive inflammasome is a complex consisting of a molecule of full-length NLRP1B, an UPA-CARD fragment, and two molecules of dipeptidyl peptidases 8 or 9 (DPP8/DPP9) ^7, 8^.

NLRP1B can be activated by direct activators that interact with the protein itself or by indirect activators. Direct activators include lethal toxin (LT) of *Bacillus anthracis*, IpaH7.8 E3 ubiquitin ligase of *Shigella flexneri*, and diverse viral proteases ^6, 9, 10^. These pathogen effectors activate NLRP1B by direct cleavage or ubiquitination of the N-terminus, resulting in proteasomal degradation up until the ZU5-UPA interface within the FIIND domain. This N-terminal degradation allows for the release of the bioactive UPA-CARD fragment, which binds caspase-1 directly to form an inflammasome ^11, 12^ through a process known as “functional degradation” ^12^. NLRP1B is also activated by the inhibitors Val-boroPro (VbP) and 1G244 ^13^, possibly by weakening the interaction between DPP8/DPP9 and NLRP1B ^7, 8^. Additional activators of NLRP1B include intracellular bacterial infections that deplete ATP ^14^, *Toxoplasma gondii* infections ^15, 16^, *Tritrichomonas musculis* infections ^17^, and agents that induce reductive stress^6, 9, 10, 18^.

It is well established that pattern-recognition receptor (PRR) signalling in intestinal epithelial cells (IECs) is important for maintaining barrier integrity and regulating immune homeostasis ^19^. Surprisingly, the roles of human NLRP1 and murine NLRP1B have not been thoroughly investigated in the intestine. In this study we made use of intestinal organoids, which recapitulate the architecture and cellular landscape of the intestinal epithelium in vitro, in order to characterise the role of NLRP1B in murine IECs.

## METHODS

### Organoid culture

Small intestinal organoids were isolated as previously described ^20–22^. Briefly, littermate WT and *Nlrp1b^-/-^* C57BL/6J mice were euthanized and ∼7 cm of the ileum was collected from each mouse and cut longitudinally. Mucus and villi were removed with a coverslip and the tissue was washed in cold phosphate-buffered saline (PBS, Wisent) and then incubated in 2 mM ethylenediaminetetraacetic acid (EDTA, Bioshop) at 4°C for 30 minutes. For generation of colonic organoids, the entire colon was collected, washed and incubated in 2 mM EDTA at 37°C for 30 minutes. Crypts were isolated by vigorous shaking and filtration though a 70-µm filter. After several washes to remove debris, crypts were resuspended in 30 µL Cultrex^TM^ Basement Membrane Extract (R&D Systems) and cultured in a 24-well plate. Small intestinal organoids were grown in Advanced Dulbecco’s modified eagle medium (DMEM)/F12 (Gibco) supplemented with 10% R-spondin-conditioned medium and 10% Noggin-conditioned medium. Colonic organoids were cultured with Advanced Dulbecco’s modified eagle medium (DMEM)/F12 (Gibco) supplemented with 20% fetal bovine serum (FBS, Wisent) and 50% L-WRN (Wnt, R-spondin and Noggin) conditioned medium ^23^. All organoid media additionally contained 2% B27 (Gibco), 1% N2 (Gibco), 1% HEPES (Gibco), 1% penicillin-streptomycin (Wisent), 1% Glutamax™ (Gibco), 500 µM N-acetylcysteine (Sigma-Aldrich), 50 ng/mL mouse epidermal growth factor (Thermo Fisher), 100 µg/mL Primocin™ (Cederlane), and 10 µM Y-27632 dihydrochloride (Tocris) for 3 days post-generation. Media was refreshed on days 3 and 5 post-seeding without Y-27632 dihydrochloride. Organoids were passaged weekly through mechanical disruption and grown in a 5% CO2 incubator at 37°C. Small intestinal organoids derived from BALB/cJ mice were cultured identically except 500 ng/mL recombinant murine R-spondin (R&D Systems) and 100 ng/mL recombinant murine Noggin (Peprotech) were used instead of R-spondin and Noggin conditioned media. Organoids were passaged at least once before use in experiments. Treatments were administered on day 3 post-passage.

### Bone marrow-derived macrophage (BMDM) culture

BMDMs were isolated from lower limbs and cultured as previously described ^24^. BMDMs were seeded at a density of 1.5×10^6^ cells per well and Cultrex^TM^ BME-seeded BMDMs were resuspended in 30 µL Cultrex^TM^ BME at 1.5×10^6^ cells per well. BMDMs were cultured in a 5% CO2 incubator at 37°C and supplemented with RPMI 1640 medium for 4 days prior to treatment.

### RAW 264.7 macrophage culture

RAW264.7 cells were purchased from ATCC (Cat.#TIB-71; RRID:CVCL_0493) and were maintained in DMEM supplemented with 10% FBS and 1% penicillin-streptomycin (Wisent) in a 5% CO2 incubator at 37°C. WT and *Nlrp1b^-/-^* RAW 264.7 macrophages were generated as previously described ^14^.

### Mice

B6.129S6-*Nlrp1b^tm1Bhk^*/J (which lack *Nlrp1a*, *Nlrp1b* and *Nlrp1c* alleles) and wild-type C57BL/6J animals were purchased from Jackson Laboratory and interbred to generate founding B6.129S6- *Nlrp1b^tm1Bhk^*/J^+/-^ heterozygotic F1 breeding pairs, hereon denoted as *Nlrp1b^+/-^* mice. All experiments were performed on F2 littermates, using corresponding WT or *Nlrp1b^-/-^* C57BL/6J pairs. Wild-type BALB/cJ animals were originally sourced from Jackson Laboratory and experimental animals were bred inhouse. All animals were maintained in a specific pathogen-free facility. Animal studies were conducted under protocols approved by University of Toronto Committee on Use and Care of Animals. Experiments were planned and conducted in a way to utilize the fewest number of animals possible. Euthanasia was performed by cervical dislocation or under deep sedation with cardiac puncture to prevent unnecessary pain or distress. Environmental enrichment was provided to all animals.

### *Tritrichomonas muris* infection and isolation

Cecal contents of *Tritrichomonas^+^* mice were collected and suspended in sterile PBS (Wisent), double filtered through 70 μM strainers (Fisherbrand), pelleted by centrifugation, repeatedly washed in PBS and then resuspended in 40% Percoll (Sigma). Cecal contents were then overlayed onto 80% Percoll and the enriched *Tritrichomonas* interphase fraction was isolated by centrifugation and extracted. Purified *Tritrichomonas* was further washed, and then filtered through a 40 μM strainer (Fisherbrand). *Tritrichomonas* were sorted based on size, granularity and green or violet autofluorescence on a FACSAriaII, as described previously ^25^. Six to eight week old WT and *Nlrp1b^-/-^* mice were orally gavaged with 2×10^6^ protozoa. Infections progressed for 21 days. At sacrifice, *Tritrichomonas* colonization was confirmed in all experimental animals via cecal content wet mount microscopy.

### Western blots

Organoids were isolated using organoid harvesting solution (R&D Systems). Dissolved Cultrex^TM^ BME was aspirated, and pellets were resuspended in 75 µL radioimmunoprecipitation assay (RIPA) buffer supplemented with 100x protease inhibitor cocktail (Sigma-Aldrich) and lysed through sonication. BMDMs (Cultrex^TM^ BME) and RAW 264.7 cells were collected in PBS and lysed as above. For whole tissue immunoblots, 0.5 cm of the distal small intestine was collected, washed of luminal content, and flash frozen in liquid nitrogen. Frozen samples were directly resuspended in 300 µL RIPA buffer supplemented with 100x protease inhibitor cocktail and 100x phosphatase inhibitor (Thermo Fisher) and sonicated on ice until visibly lysed.

Protein concentrations were determined by bicinchoninic acid assay (Thermo Fisher) using the manufacturer’s protocol. Lysates were run on SDS-PAGE using standard techniques. Gels were transferred onto polyvinylidene fluoride membrane using a semi-dry transfer apparatus (Trans-Blot Turbo Transfer System, Bio-Rad) and were blocked for 45 minutes in 5% bovine serum albumin (BSA, Sigma-Aldrich) in tris-buffered saline and 0.1 Tween 20 (TBST), before addition of primary antibodies. Primary and secondary antibodies were incubated overnight with rotation at 4°C, or at room temperature for 1 hour, respectively. Membranes were incubated with Luminata Crescendo or Classico Western Horseradish Peroxidase Substrate (Millipore Sigma) and visualized using a G:Box Chemi XX6 (Syngene).

### In situ hybridization

For *in situ* hybridization, FFPE tissue was sectioned (5 μM), deparaffinized and rehydrated according to manufactures instructions. Tissue was hybridized and amplified using the RNAscope 2.5 HD assay (Advanced Cell Diagnostics, ACD, 322370) and all probes were sourced from ACD. A minimum of two biological repeats per probe were used. For each experiment, negative and positive control probes, supplied by ACD were run in parallel with target probes.

### Antibodies

α/β-Tubulin (1:1000, WB) (Cell-Signaling Technology, 2148), Cleaved caspase-3 (CST, 9661S), Cleaved caspase-8 (CST, 8592S), IL-18 (1:1000, WB) (Abcam, ab207323), GSDMC2/GSDMC3 (Abcam, ab229896), GSDMC3 (Abclonal, A16741), GSDMD (1:1000, WB) (Abcam, ab209845), GSDME (Abcam, ab215191), NLRP1B (1:1000, WB) (Adipogen, AG-20B-0084-C100), caspase-1 (1:1000, WB) (Adipogen, AG-20B-0042-C100), MEK1 (1:1000, WB) (Sigma-Aldrich, 07-641), DPP9 (1:1000, WB) (Abcam, ab302903).

### Quantitative polymerase chain reaction (qPCR)

For organoids, BMDMs and cell lines, RNA extraction, DNAse treatment, and cDNA synthesis, were all performed using the GeneJET RNA purification kit (Thermo Fisher), TURBO DNA-free kit (Invitrogen), and the high-capacity cDNA reverse transcription kit (Applied Biosystems) according to manufacturer’s protocols. For whole tissue samples, RNA was extracted using the Trizol^TM^ Reagent according to manufacturer’s protocols. The resulting cDNA was diluted accordingly before qPCR with PowerUp^TM^ SYBR^TM^ Green Master Mix (Applied Biosystems ThermoFisher). Cycle threshold (Ct) values were acquired from a CFX384 TouchTM Real-Time PCR Detection System (Bio-Rad) with 40 cycles of 60°C annealing/extension for 1 minute, followed by a melt curve. Gene expression was normalized to the average of two housekeeping genes (*Gapdh*, *Rpl19*). Primers were designed using NCBI primer blast with the following criteria: annealing temperature of 60°C, maximal product length of 200bp, exohe n-exon spanning, and GC content of approximately 50%. Primer sequences are listed below. Statistics were performed with the Student’s t-test or Two-tailed ANOVA with Tukey’s multiple comparisons test, where appropriate.

### qPCR primers (sequences are 5’ to 3’)

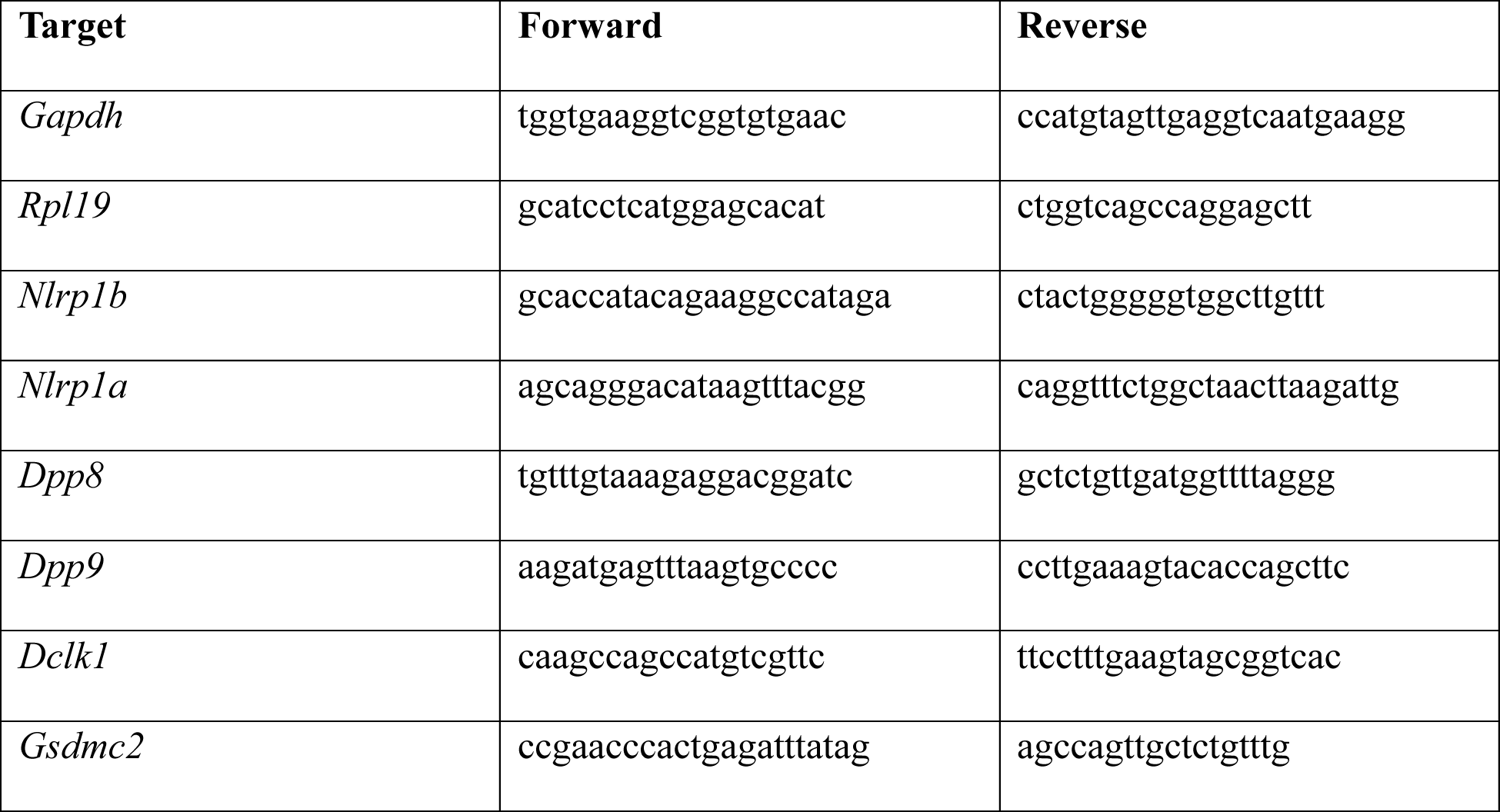

### Fluorescent Dpp substrate assay

Organoids were seeded in 40 μL Cultrex^TM^ BME in a 96-well plate and grown in 200 μL organoid media replaced daily. Stimulated organoids were incubated with the fluorogenic DPP substrate 1 Ala-Pro-AMC dipeptide (BPS Bioscience), at 1:20 in PBS. Fluorescence was measured after 10 minutes by SpectraMax®i3x (excitation 350 nm, emission 450 nm) and reported in relative fluorescence units (RFUs). An unpaired t test was used to detect significance between groups.

### Chemicals and reagents

1G244 (Millipore Sigma), 5z7 (Sigma-Aldrich), LF (List Labs), Ifn-γ recombinant mouse protein (Thermo Fisher), recombinant mouse IL-22 protein (R&D Systems), recombinant murine IL-13 (Peprotech), recombinant murine TNF-α (Peprotech), standard flagellin from *Salmonella typhimurium* (Invivogen), and Val-Boro-Pro (Sigma-Aldrich). PA was purified using previously described methods ^26^.

## RESULTS

### *Nlrp1b* is expressed in the small intestinal epithelium of C57BL/6J mice

Bulk RNA-sequencing data generated for an ongoing study in our laboratory (Ranger et al., manuscript in preparation) revealed that *Nlrp1b* gene expression was present at baseline in wild-type (WT) murine ileal organoids from C57BL/6J mice (Fig. 1A). Relative expression levels defined by reads per kilobase of transcript per million reads mapped (RPKM) showed that *Nlrp1b* expression was lower than some other NLRs, including *Nlrp6* and *Nlrc4*, which have been shown to have functionality in the intestinal epithelium ^27–32^, but higher than *Nod2 and Nlrc3* (Fig. 1A). In addition to *Nlrp1b*, some mice express the paralog *Nlrp1a*, which like *Nlrp1b* contains all domains associated with functioning murine NLRs ^33^. In contrast, the mouse paralog *Nlrp1c* lacks a CARD domain and is currently classed as a pseudogene ^33^. Notably, we observed no detectable *Nlrp1a* expression in C57BL/6J ileal organoids (Fig. 1A). To confirm *Nlrp1b* expression, we conducted quantitative RT-PCR (qPCR) and RNAscope^TM^ in situ hybridization analyses. For the qPCR assays, both ileal organoids and bone marrow-derived macrophages (BMDMs) were prepared from the same WT C57BL/6J mice, and baseline *Nlrp1b* and *Nlrp1a* expression was compared between the two samples. *Nlrp1b* mRNA was detected in SI organoids, but at a lower level than was found in BMDMs (Fig. 1B), whereas *Nlrp1a* mRNA was only detectable in BMDMs (Fig. 1C), suggesting that *Nlrp1b* is the dominant paralog expressed in the C57BL/6J intestinal epithelium. Next, we queried the spatial localization of *Nlrp1b* transcripts in ileal tissue from WT C57BL/6J mice and observed weak expression in the mid- to upper-villus region (Fig.1D and data not shown). Because the visualization of *Nlrp1b* transcripts was at the limit of detection of the RNAscope^TM^ assay, we used an orthogonal approach to determine the IEC sub-populations in which *Nlrp1b* transcripts were expressed. We extracted data from a single cell RNAseq analysis of ileal organoids from WT C57BL/6J mice performed for another project (Goyal et al., manuscript submitted). We observed that *Nlrp1b* transcripts were expressed at low levels in IECs without an obvious tropism for a specific IEC subset, although expression was slightly higher in the most mature enterocytes (Fig. 1E, arrow) marked by expression of *Slc2a2* and *Ada* (previously identified as mid- and upper-villi markers, respectively ^34^), supporting our RNAscope^TM^ data showing expression in villi enterocytes. To determine whether NLRP1B protein expression could be detected in murine SI organoids, we used a commercial antibody that targets the C-terminal CARD of NLRP1B and validated it using BMDM lysates from WT and *Nlrp1b^-/-^* C57BL/6J mice. Using Western blotting, we observed specific bands at molecular weights ∼25 kDa and ∼134 kDa, corresponding to the UPA-CARD fragment and the full-length unprocessed NLRP1B, respectively (Figs. 1G and 1F). Interestingly, NLRP1B was found at much greater levels in IECs than BMDMs, despite seeing the inverse correlation at the mRNA level (Fig. 1H), and was mainly detected as the UPA-CARD fragment. Moreover, NLRP1B protein levels were similar in SI and colonic organoids (Fig. 1H, compare lanes 1 and 5). As a positive control, lysates from RAW264.7 murine macrophages were used, which showed very strong expression of both the full-length and UPA-CARD forms (Fig. 1H, lane 6). We note that while RAW264.7 macrophages express allele 1 of *Nlrp1b* and C57BL/6J mice express allele 2, the CARD domains of the two proteins that the antibody recognizes have identical protein sequences. Caspase-1 was expressed at comparable levels in C57BL/6J BMDMs, SI and colonic organoids, while the caspase-1 substrates GSDMD and IL-18 were expressed more strongly in SI and colonic organoids than in BMDMs (Fig. 1H). Together, these results indicate that the proteins required for NLRP1B-dependent pyroptosis are present in ileal and colonic IECs.

**Figure 1:**
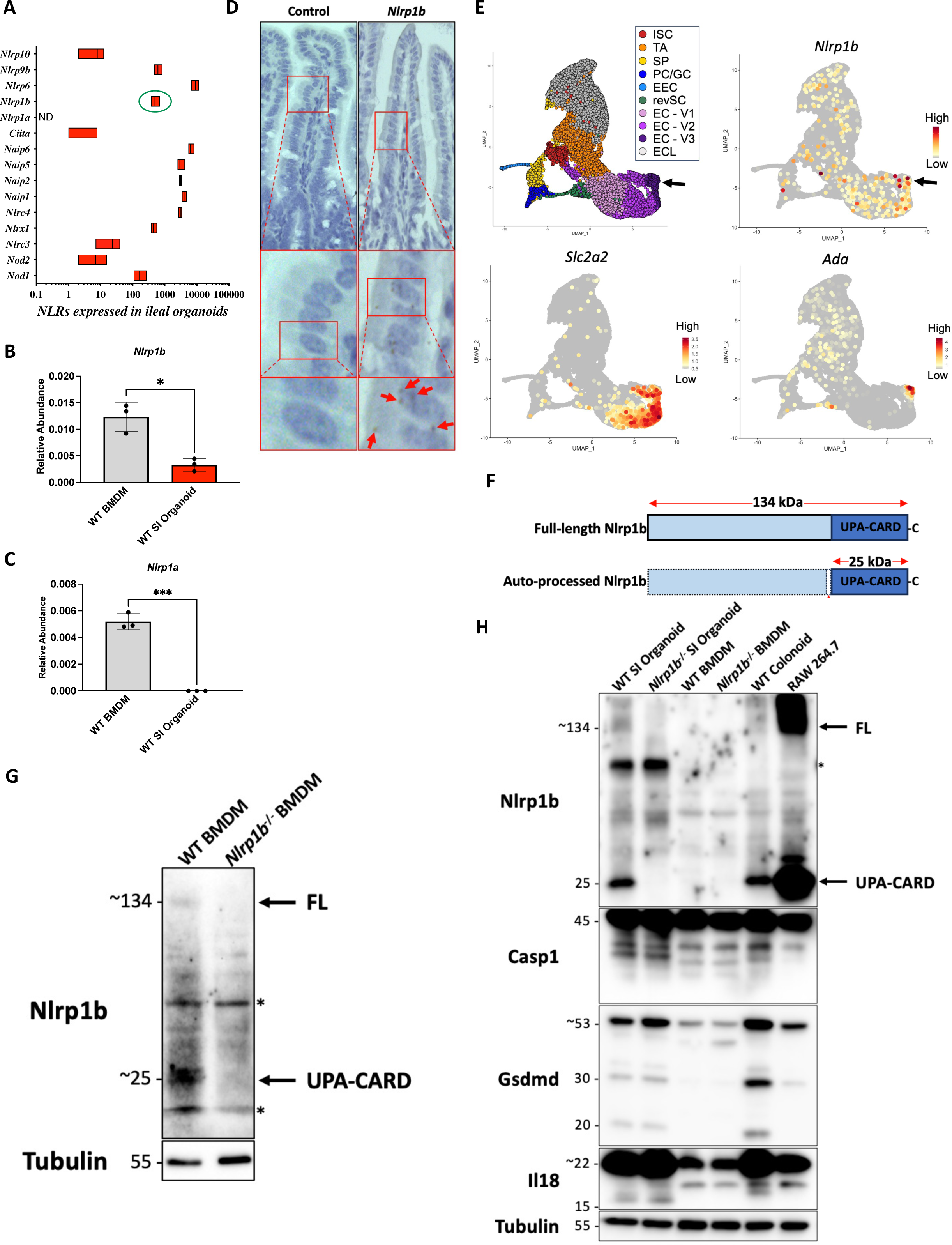
NLRP1B is expressed in the small intestinal epithelium of C57BL/6J mice. **(A)** RNA-sequencing analysis of NLRs expressed in WT ileal organoids. X-axis represents reads per kilobase of transcript per million reads mapped (RPKM). **(B** and **C)** qPCR analysis of *Nlrp1b* expression in WT BMDMs and WT murine ileal organoids. **(D)** RNAscope^TM^ *in-situ* hybridization assay probing for *Nlrp1b* and the negative control target, *DapB* from *Bacillus subtilis*, in the ileum of WT C57BL/6J mice. **(E)** UMAP plot of single cell RNAseq data set from murine derived ileal organoids. Cell types are annotated as followed: intestinal stem cells (ISC), transit amplifying cells (TA), secretory progenitor (SP), goblet and Paneth cells (GC/PC), enteroendocrine cells (EEC), revival stem cells (revSC), enterocytes (EC–V1/V2/V3) and enterocyst-like cells (ECL). Expression of *Nlrp1b, Slc2a2* and *Ada* overlayed on UMAP plot. Arrow indicates the V3 region of the villus enterocytes corresponding to the upper villi (where Ada expression is maximal) in which Nlrp1b was the greatest. **(F)** Schematic illustration of full length NLRP1B, its auto-processed form, and UPA-CARD domain. **(G)** Validation of NLRP1B antibody by Western blot with protein lysates from WT and *Nlrp1b*^-/-^ BMDMs. Tubulin was used as a loading control. (**H**) Western blot analysis of baseline NLRP1B expression level in WT and *Nlrp1b*^-/-^ organoids and BMDMs. Protein lysates were probed for inflammasome components NLRP1B, caspase-1, GSDMD, and IL-18.

### VbP does not activate NLRP1B in SI organoids derived from C57BL/6J mice

We used Val-boro-Pro (VbP) to examine the function of NLRP1B in IECs from C57BL/6J SI organoids. VbP is a dipeptidyl peptidase inhibitor which can activate human and rat NLRP1 as well as all functional murine *Nlrp1b* alleles, including alleles 1 and 2 ^13^. To our surprise, VbP did not activate NLRP1B in SI organoids from C57BL/6J mice, as evidenced by a lack of cleavage of GSDMD and Caspase-1 (Fig. 2A). This was not because SI organoids were refractory to caspase-1 activation since bacterial flagellin, a specific ligand of the NAIP/NLRC4 inflammasome, potently induced GSDMD and Caspase-1 cleavage in an NLRP1B-independent manner (Fig. 2A). To account for a potential inhibitory effect on VbP of the Cultrex^TM^ basement membrane extract (BME), which is required for intestinal culture, we plated BMDMs in an equivalent volume of BME as was used for organoids. Despite the fact that NLRP1B protein levels were relatively low in C57BL/6J BMDMs as compared to SI organoids (Fig. 2B, compare lanes 1-3 with 7-8; see also above Fig. 1H), the BMDMs, but not the organoids, responded to VbP, leading to downstream GSDMD cleavage in an NLRP1B-dependent manner (Fig. 2B). Next, we asked if organoids had a lower sensitivity or delayed response to VbP. Increasing the concentration of VbP from 10 μM to 100 μM and extending the duration of SI organoid treatment from 4 h to up to 72 h had no effect, however, on NLRP1B inflammasome activation (Fig. 2C). A number of agents synergize with VbP to induce greater NLRP1/NLRP1B activation, including the antioxidant JSH-23, which accelerates N-terminal degradation of NLRP1/NLRP1B ^18^. However, treatment of wild-type organoids with JSH-23 in combination with VbP did not result in NLRP1B inflammasome activation, as assessed by GSDMD cleavage (data not shown).

**Figure 2:**
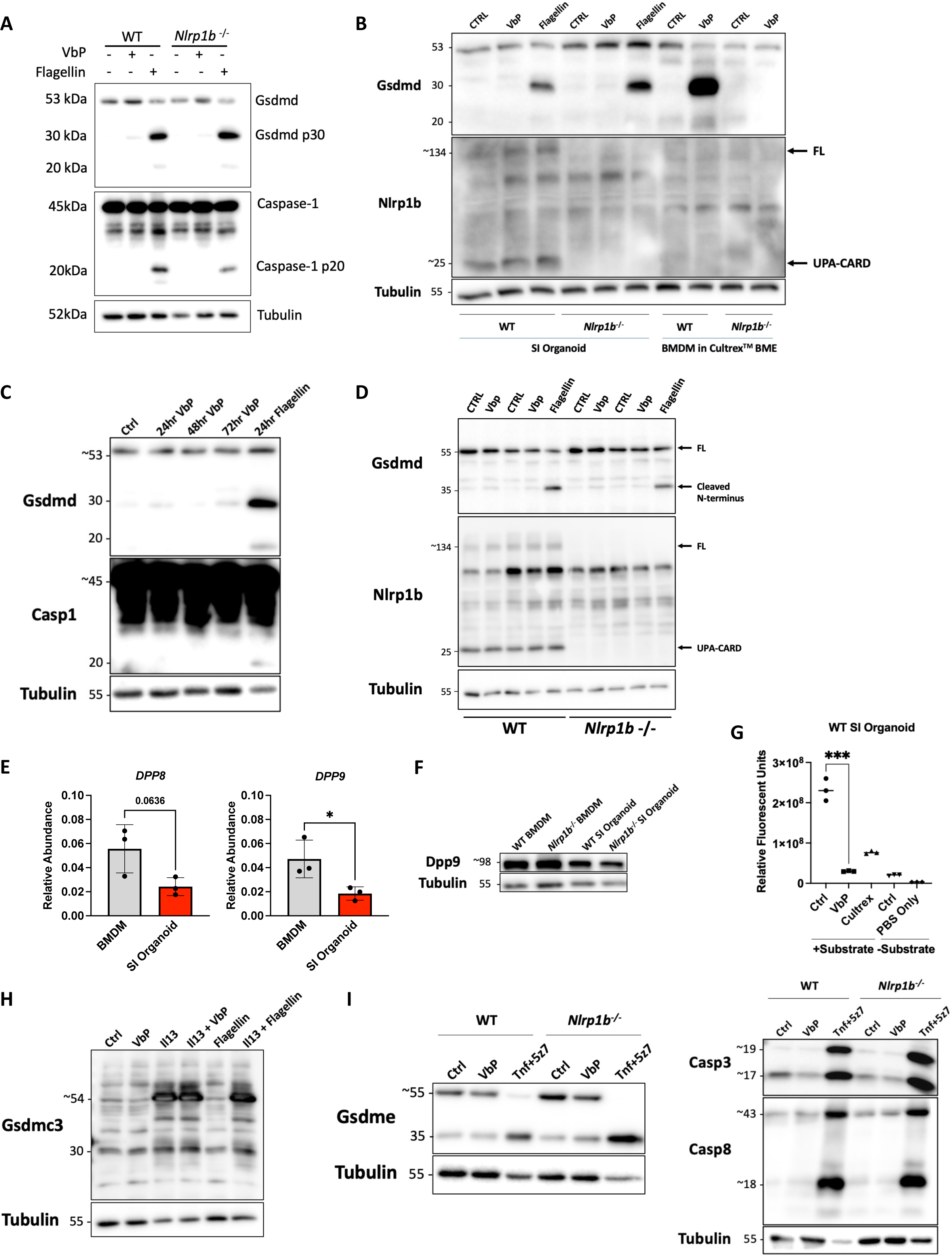
VbP does not induce downstream Gsdmd cleavage or other pyroptotic pathways in SI organoids from C57BL/6J mice. **(A)** Murine WT and *Nlrp1b*^-/-^ SI organoids were treated with VbP (10μM) or flagellin (2 μg/mL) for 4 hours. **(B)** SI organoids and BMDMs derived from WT and *Nlrp1b*^-/-^ mice were treated with VbP (10 μM) or flagellin (2 μg/mL) for 24 hours; both seeded in 30 μL Cultrex^TM^ BME. The same experiment was done for longer timepoints in WT SI organoids in **(C)**. Time points for VbP were 24, 48, or 72 hours, and 24 hours for flagellin. **(D)** WT and *Nlrp1b*^-/-^ SI organoids were treated with larger dose of VbP (100μM) or flagellin (5 μg/mL) 3 to 4 days post passaging. **(E)** qPCR analysis of *Dpp8* and *Dpp9* in WT C57BL/6J BMDMs and SI organoids. **(F)** Western blot analysis of baseline expression of DPP9 protein in WT and *Nlrp1b*^-/-^ BMDMs and SI organoids. **(G)** Statistical analysis of relative fluorescence units (RFUs) measured by SpectraMax^®^i3x (excitation 350 nm, emission 450 nm) in WT SI organoids after 1 hour incubation with Ala-Pro-AMC dipeptide, followed by treatment of VbP (100 μM). \*\*\**P* < 0.001. **(H)** WT SI organoids were primed with IL-13 for 24 hours, then treated with VbP (100 μM) or flagellin (2 μg/mL) for 24 hours. **(I)** WT and *Nlrp1b*^-/-^ SI organoids were treated with 4 hours of VbP (100 μM) or TNF-α (10ng/ml) and 5z7 (2.5μM).

Organoids become more differentiated as a function of time. Therefore, we wondered if NLRP1B was expressed only in a transient IEC population. However, stimulation at different stages of organoid growth (day 2 up to day 5 post-passaging) (Fig. 2D and data not shown) had no effect on GSDMD cleavage, showing that VbP was unable to trigger inflammasome activation through NLRP1B in C57BL/6J SI organoids, in contrast to C57BL/6J BMDMs. In addition to GSDMD, we also assessed Caspase-1 cleavage and found that while flagellin could trigger the formation of the p20 form of Caspase-1, VbP was unable to do so (Fig. 2C).

We next aimed to determine whether this discordance between NLRP1B protein expression and functionality was due to increased DPP8 and DPP9 levels in organoids. Surprisingly, small intestinal organoids displayed decreased levels of *Dpp9* transcripts (Fig. 2E) and protein expression (Fig. 2F) when compared to BMDMs. Likewise, *Dpp8* transcripts were similarly reduced (p=0.06) in the organoids (Fig. 2E). Further, to ensure that VbP effectively inhibited organoid DPPs, we used a fluorogenic assay that measures the fluorescence of the fluorophore, AMC, which is released following the DPP-mediated cleavage of Ala-Pro-AMC. We found that VbP significantly reduced fluorescence, indicating that it reduced DPP activity in organoids (Fig. 2G). Collectively, these results suggest that SI organoids were not refractory to VbP-mediated effects because of greater levels of DPP expression, or lack of VbP-driven inhibition of DPPs.

While VbP was unable to trigger GSDMD and caspase-1 cleavage in SI organoids, we reasoned that NLRP1B could possibly activate a non-canonical inflammasome pathway. Therefore, we also used Western blotting to assess the cleavage of GSDMC and GSDME, induced by Caspase-8 and Caspase-3, respectively ^35–38^. Since GSDMC expression has been shown to be massively induced by both enteric parasites and cytokines involved in the anti-parasitic response in the SI ^39, 40^, such as IL-4/IL-13, we also stimulated SI organoids with IL-13 in order to upregulate GSDMC expression. However, VbP was unable to induce cleavage of GSDMC, at baseline or following upregulation by IL-13 treatment (Fig. 2H), and was also unable to cleave GSDME, while the positive control treatment of TNF-α + 5z7 potently induced GSDME cleavage (Fig. 2I, left), as well as Caspase 3/8 activation (Fig. 2I, right). Thus, we conclude that while NLRP1B was expressed in C57BL/6J SI organoids, IECs remained refractory to VbP-dependent activation of the NLRP1B inflammasome.

### IL-13 stimulation and *Tritrichomonas* infection induce *Nlrp1b* expression in C57BL/6J mice

In macrophages, some NLRs require priming by MAMPs, DAMPs or cytokines to induce expression or elicit post-translational modifications required for inflammasome activation ^2, 3, 41^. Accordingly, it is plausible that NLRP1B was not activated because it requires priming in IECs. Therefore, we tested whether IFN-γ, IL-22 or IL-13, three crucial cytokines involved in the control of epithelial responses to infection and inflammation in the intestine, were required for priming NLRP1B in C57BL/6J SI organoids. While stimulation of WT C57BL/6J organoids with IFN-γ or IL-22 for 4 h, 8 h and 24 h, had no effect on *Nlrp1b* transcript levels (Fig. 3A), we found that IL-13, a cytokine produced during parasitic infection, significantly induced *Nlrp1b* at 24 h (Fig. 3B). We thus aimed to determine if IL-13 priming was required for VbP-dependent activation of NLRP1B. However, co-treatment with IL-13 and VbP did not result in NLRP1B-dependent GSDMD cleavage (Fig. 3C). Nevertheless, the moderate (2-3 fold, see Fig. 3A) induction of *Nlrp1b* transcript by IL-13 is suggestive of a potential role of this NLR protein in anti-parasitic responses in the gut, in line with a recent report which showed that *Nlrp1b^-/-^* mice had defective host responses to the protozoan *Tritrichomonas musculis* ^17^. In support of this idea, *Nlrp1b* expression was upregulated in the SI tissue of C57BL/6J mice acutely infected by oral gavage for 3 weeks with *Tritrichomonas muris*, as compared to non-infected littermate animals (Fig. 3D) and RNAscope^TM^ analysis of the SI tissue also showed stronger staining in the infected tissue, which was more obvious in the mid to upper region of the epithelial villi (Fig. 3E). Moreover, *Tritrichomonas* infection resulted in an up-regulation of NLRP1B in protein extracts from isolated SI tissue (Fig. 3F), although we acknowledge that this observation may be the consequence of the recruitment of immune cells expressing NLRP1B rather than the increased expression in the epithelium. We were unable to track the cell population(s) in which NLRP1B protein expression was increased following infection because our anti-NLRP1B antibody was not suitable for immunofluorescence (data not shown). Together, these data suggest that the previously identified ^17^ role for NLRP1B in the anti-parasitic response to *Tritrichomonas* may be, at least in part, dependent on the expression of NLRP1B in IECs, but IL-13-dependent priming does not facilitate VbP-mediated inflammasome activation in SI IECs from C57BL/6J mice.

**Figure 3:**
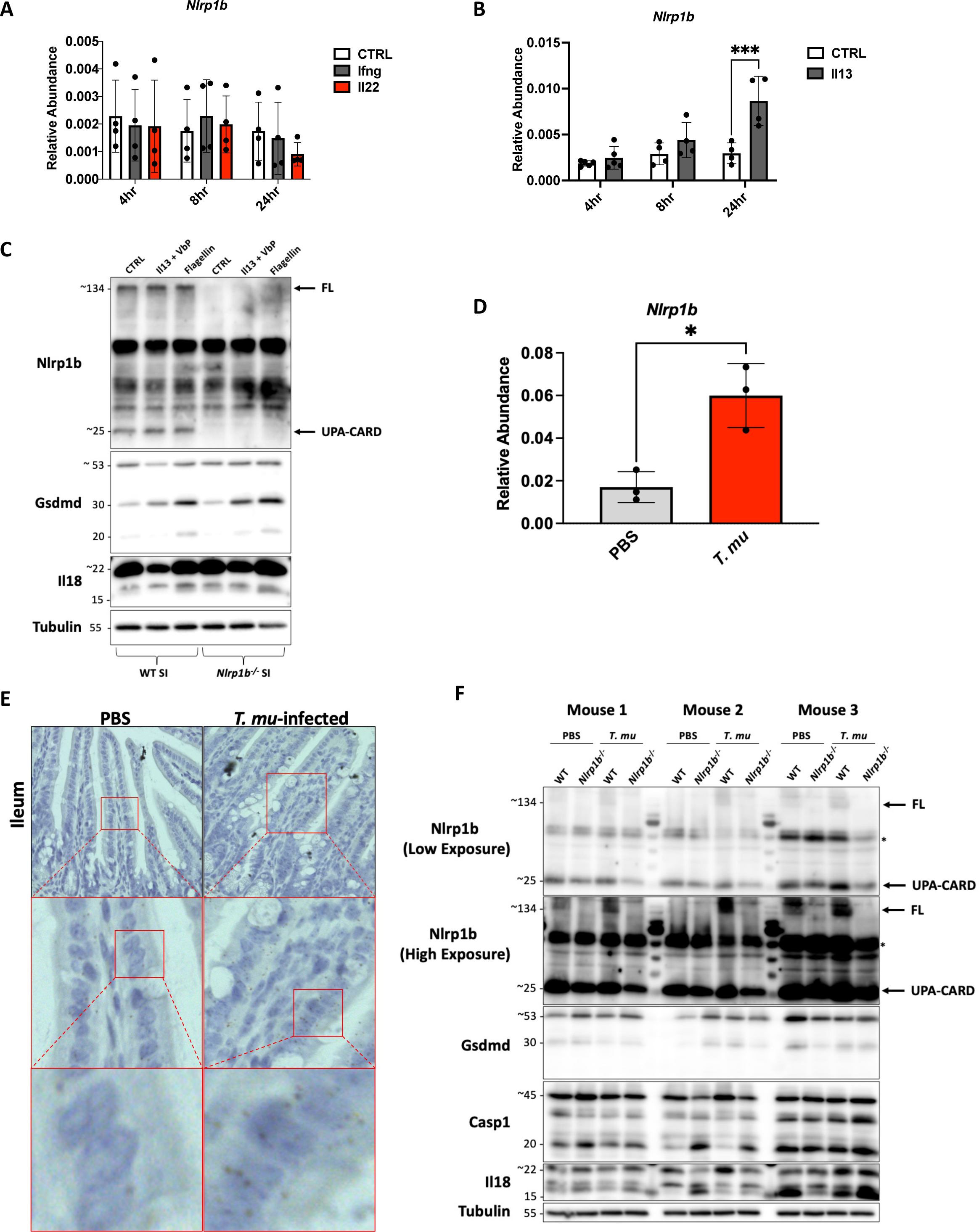
IL-13 stimulation and *Tritrichomonas* infection induce *Nlrp1b* expression. (**A and B**) qPCR expression of *Nlrp1b*. SI organoids derived from WT C57BL/6J mice were treated with IFN-ɣ (10 ng/mL), IL-22 (100 ng/mL), or IL-13 (2 ng/mL) for 4, 8 or 24 hours. \**P* < 0.05**. (C)** WT and *Nlrp1b^-/-^* C57BL/6J SI organoids were co-treated with IL-13 (2 ng/mL) and VbP (100 μM), or flagellin (2 μg/mL) for 24 h. Protein lysates were collected for Western blot analysis. **(D-F)** Littermate WT and *Nlrp1b^-/-^* mice were orally gavaged at age 6 to 8 weeks with 2×10^6^ *Tritrichomonas muris* (*T. mu*) parasites in PBS. Mice were sacrificed 21 days post-infection for ileum tissue collection. **(D)** qPCR expression of *Nlrp1b*, \*\*\**P* < 0.001. **(E)** RNAscope^TM^ *in-situ* hybridization assay probing for *Nlrp1b* in the ileum of WT and *T.mu* infected mice. **(F)** Whole tissue western blot analysis of NLRP1B, GSDMD, caspase-1, and IL-18 protein expression in ileum.

### VbP does not activate NLRP1B in colonoids from C57BL/6J mice

After assessing the functionality of NLRP1B in the SI, we next analyzed a potential role in colonic IECs. We observed earlier that WT C57BL/6J colonoids expressed NLRP1B at comparable levels to SI organoids (see Fig. 1H). However, unlike in the SI, *Nlrp1b* transcripts were not induced by IL-13 in C57BL/6J colonoids, while the cytokine potently induced *Gsdmc2,* an IL-13 target gene (Fig. 4A) ^40^. In agreement, analysis of *Nlrp1b* expression in the colon of C57BL/6J mice by RNAscope^TM^ showed that the transcript was found mainly in the upper part of the colonic crypts, but that the expression level was not different in colonic sections from *Tritrichomonas*-infected animals (Fig 4B). Additionally, we found that IL-13 did not substantially induce NLRP1B at the protein level in C57BL/6J colonoids, while GSDMC expression was greatly upregulated (Fig. 4C). Moreover, IL-13 did not prime inflammasome activation with VbP in C57BL/6J colonoids (Fig 4D). Together, these results show that while NLRP1B is expressed in colonic IECs from C57BL/6J mice, this tissue remained refractory to VbP-dependent inflammasome activation.

**Figure 4:**
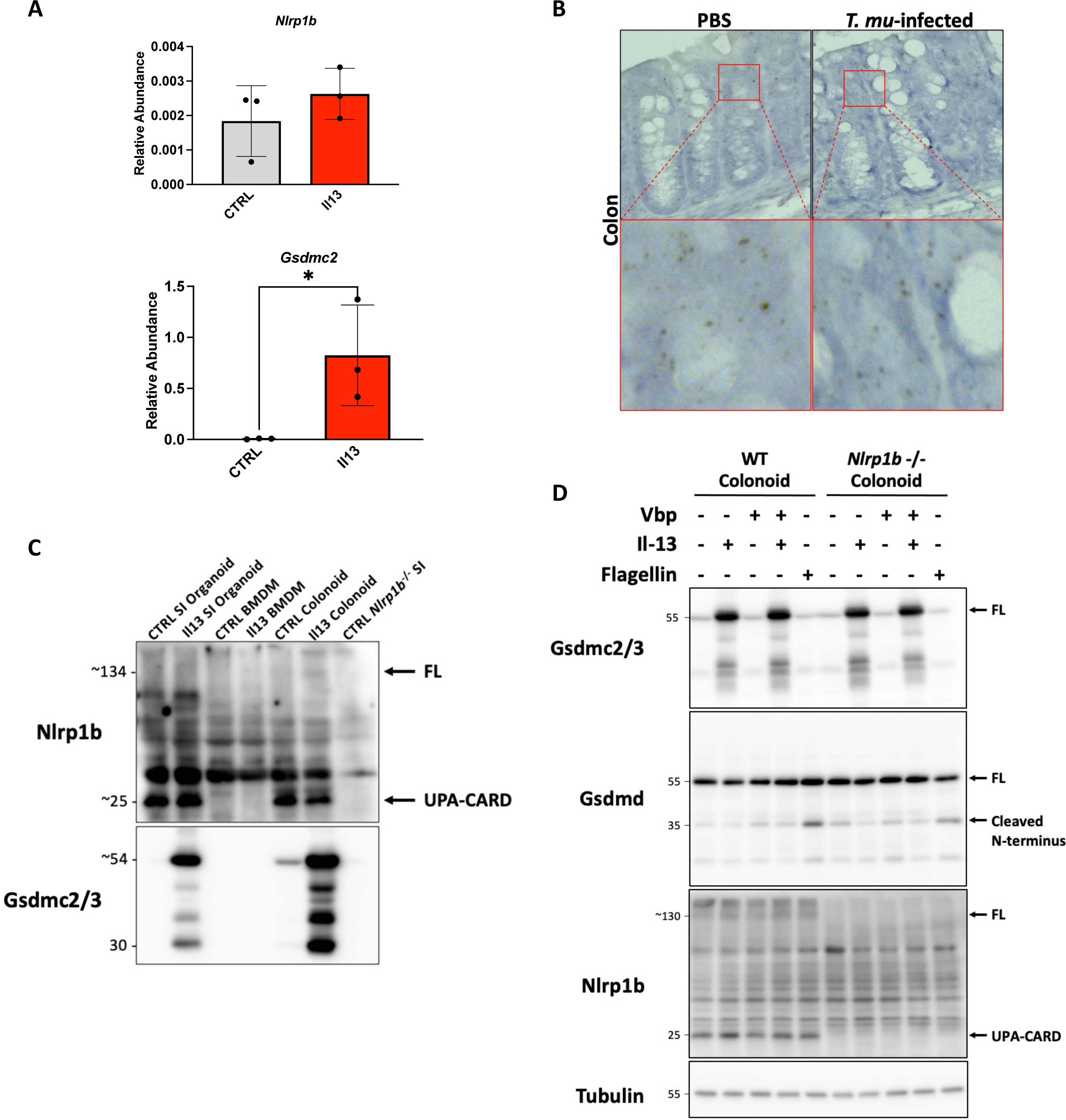
Vbp does not trigger downstream Gsdmd cleavage in colonoids from C57BL/6J mice. **(A)** qPCR expression of *Nlrp1b* and *Gsdmc2* in colonoids treated with IL-13 (2 ng/mL) for 24 hours. Colonoids were derived from WT C57BL/6J mice. CT value normalized to house-keeping genes *Gapdh* and *Rpl19.* \**P* < 0.05. **(B)** RNA Scope^TM^ *in-situ* hybridization assay probing for *Nlrp1b* in the colon of *T.mu* infected and non-infected WT C57BL/6J mice. **(C)** WT C57BL/6J SI organoids, colonic organoids, and BMDMs were treated with IL-13 (2 ng/mL) for 24 hours. **(D)** WT and *Nlrp1b^-/-^* C57BL/6J colonoids were treated with IL-13 (2 ng/mL), VbP (100μM), flagellin (5μg/ml) or IL-13 + VbP for 24 hours.

### NLRP1B is not detected in organoids derived from BALB/cJ mice

While we found that NLRP1B encoded by allele 2 in C57BL/6J mice was refractory to activation in IECs, we aimed to test whether NLRP1B in organoids derived from BALB/cJ mice expressing allele 1 could be activated. To our surprise, we found that *Nlrp1b* expression was undetectable by qPCR in SI organoids from BALB/cJ mice (Fig. 5A). This was not a result of allelic specificity in our PCR primers as they targeted a common region between the two alleles of *Nlrp1*; moreover, we could amplify *Nlrp1b* allele 1 transcripts from RAW264.7 macrophages (data not shown). NLRP1B was not detectable by Western blotting in lysates from BALB/cJ SI organoids (Fig. 5B, lane 3), but was readily detectable in BALB/cJ BMDMs (Fig. 5B, lane 6). Although NLRP1B was not detected in BALB/cJ SI organoids, we reasoned that trace levels could still be functional. Therefore, we assessed whether we could activate NLRP1B in BALB/cJ SI organoids by using VbP, the specific DPP8/DPP9 inhibitor 1G244, and anthrax Lethal Toxin (LT), a robust and well-studied activator of NLRP1B encoded by allele 1 but not allele 2 ^42^. As a control, we used WT and *Nlrp1b^-/-^* RAW 264.7 cells to confirm that VbP, 1G244, and LT induced inflammasome activation in an NLRP1B-dependent manner (Fig. 5C). As expected, LT was unable to activate NLRP1B-dependent GSDMD cleavage in C57BL/6J organoids because these cells express *Nlrp1b* allele 2 (Fig. 5D). In BALB/cJ SI organoids, LT was also unable to induce GSDMD cleavage (Fig. 5D), and this was not a consequence of LT being unable to enter the cytosol in these organoids since LT treatment induced a reduction in the levels of MEK1, a protein known to be a direct target of the LF protease subunit of LT (Fig. 5D) ^43^. In addition, VbP and 1G244 were also unable to induce GSDMD cleavage in BALB/cJ SI organoids (Fig. 5D). We therefore conclude that BALB/cJ-derived SI organoids do not express a functional NLRP1B inflammasome.

**Figure 5:**
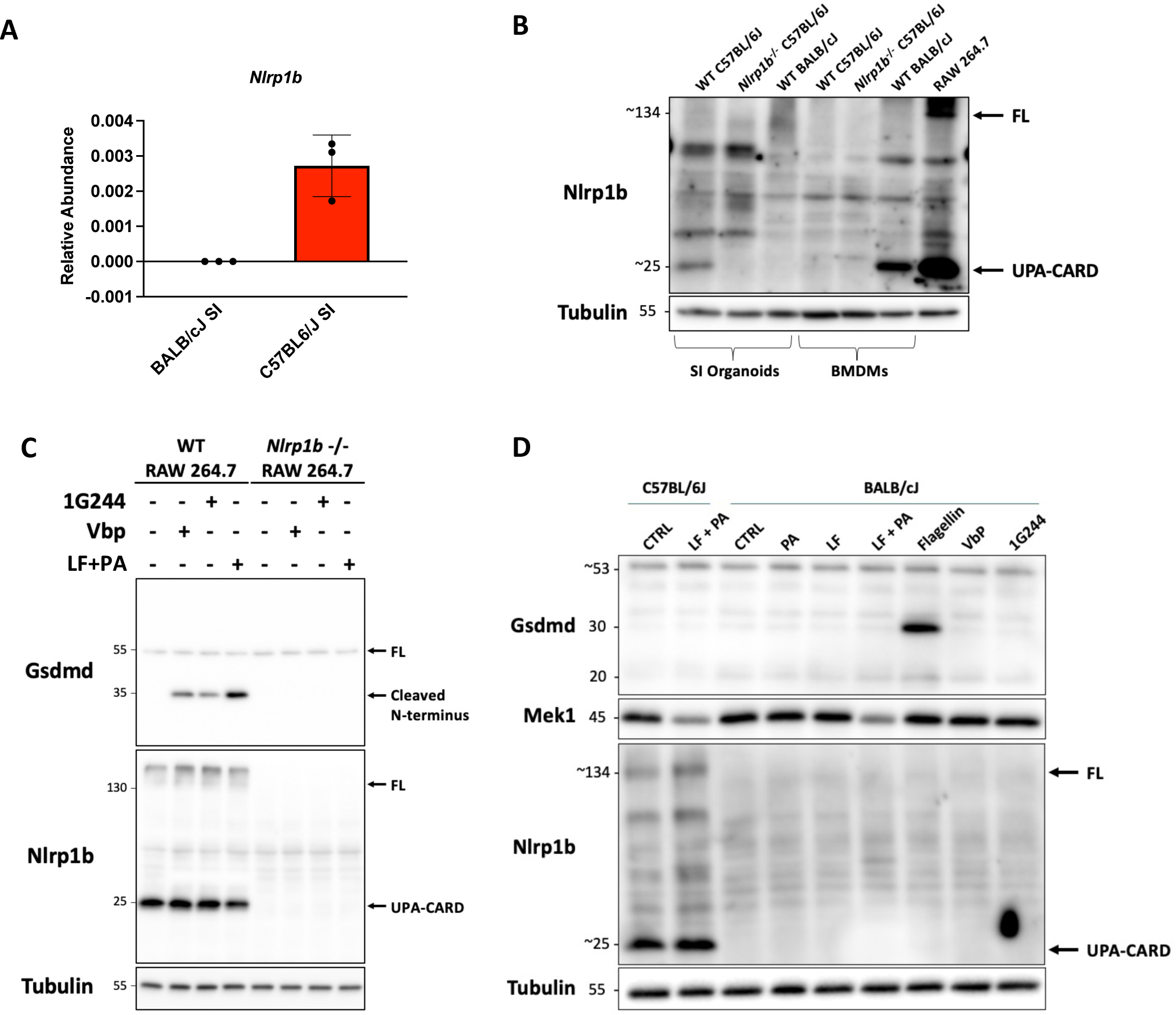
NLRP1B is not expressed in organoids from BALB/cJ mice. **(A)** qPCR expression of baseline *Nlrp1b* in WT BALB/cJ and C57BL/6J SI organoids. **(B)** Western blot analysis of NLRP1B protein expression in untreated SI organoids and BMDMs derived from WT and *Nlrp1b^-/-^* C57BL/6J mice, WT BALB/cJ mice, and RAW 264.7 macrophages. **(C)** WT and *Nlrp1b*^-/-^ RAW 264.7 macrophages were treated with VbP (10 μM) or 1G244 (10 μM) or Lethal Toxin [LF (10 nM) with PA (1 nM)] for 3 hours. **(D)** Western blot analysis of GSDMD, MEK1, and NLRP1B protein expression in WT C57BL/6J or BALB/cJ SI organoids. C57BL/6J organoids were treated with LF (20 nM) + PA (2 nM) for 4 h. BALB/cJ organoids were stimulated with LF (20nM), PA (2nM), LF (20nM) + PA (2nM), flagellin (10 μg/mL), VbP (10 μM), or 1G244 (10 μM) for 4 hours.

## DISCUSSION

In this work, we have provided the first characterization of the NLRP1B inflammasome in IECs of the murine intestinal epithelium. We found that *Nlrp1b* mRNA transcripts were detectable in murine colonic and ileal organoids and corresponding whole intestinal tissue. RNA in situ hybridization revealed that *Nlrp1b* expression was not limited to a specific IEC cell-type, although mid to upper-villi enterocytes exhibited more pronounced staining, which we confirmed by analyzing scRNAseq data from murine SI organoids.

To our surprise, a greater amount of NLRP1B protein was detected in C57BL/6J ileal organoids than in BMDMs derived from the same animals. This is in contrast to our qPCR data, which showed higher *Nlrp1b* mRNA levels in BMDMs, suggesting that NLRP1B may undergo tissue-specific post-transcriptional regulation. Despite the difference in protein levels, VbP was capable of activating NLRP1B only in BMDMs. This was not a result of a failure of VbP to inhibit DPPs as confirmed via fluorogenic Ala-Pro-AMC assay. However, one caveat of this assay in organoids is that the inhibition of Ala-Pro-AMC cleavage observed may be mainly driven by DPP4 inhibition, which is plasma membrane-bound in IECs ^44^. To compensate for this, we increased the VbP dosage 10-fold, increased the duration of treatment, and also tested a DPP8/DPP9-specific inhibitor, 1G244, all to no avail.

Notably, intestinal organoids appear to have reduced expression of DPP8 and DPP9 when compared to BMDMs generated from the same C57BL/6J animals. Given that VbP is thought to activate NLRP1B by inhibiting DPPs ^7, 8^, one would expect that reduced DPP expression, coupled with elevated NLRP1B levels would prime the epithelium for inflammasome activation. One possibility is that DPP expression is tightly regulated in order to maintain intestinal homeostasis. For example, DPP4 is a receptor for MERS-CoV infection ^45^, and elevated serum DPP4 levels have been associated with numerous viral infections, including hepatitis C virus and Epstein-Barr virus^46^. Likewise, increased fecal DPP4 has recently been proposed as a biomarker for ileal Crohn’s disease ^47^. While less is known about the function of DPP8 and DPP9 in the intestine, transcriptome analysis has linked elevated intestinal DPP9 expression with genes relating to viral replication ^48^, while genome wide association studies have connected high DPP9 expression to worsened COVID-19 pathology ^49^. As NLRP1 can be activated by viral proteases ^50^ it is possible that in the murine epithelium, NLRP1B and DPP8/9 may be involved in anti-viral responses, and could be held under regulatory mechanisms to prevent against aberrant intestinal inflammation.

A known repressor of the NLRP1B inflammasome is an oxidized cell state. Specifically, activation is restrained by the oxidized form of thioredoxin-1 (TRX1), which is associated with the NACHT-LRR region of NLRP1B and represses N-terminal degradation during oxidative stress ^18, 51, 52^. While intestinal inflammation is a source of oxidative stress for IECs ^53^, all intestinal organoids used in this study were cultured under reducing conditions. Organoid growth media is composed of serum analogs such as B27 that contains reduced glutathione ^54^, and which has been shown to decrease binding NLRP1 to TRX1 ^18, 51, 52^. Likewise, sodium pyruvate, which is protective against H2O2-induced oxidative stress ^55^, and N-acetyl-cysteine (NAC), which can directly scavenge ROS species and as a precursor to glutathione, can also be converted to neutralize ROS species during oxidative stress ^56^, are also present in our cultures. While it remains to be tested whether BMDMs are less oxidized than IECs, the presence of numerous reducing agents in our culture conditions suggests that oxidative stress is unlikely to be the factor contributing to the refractive state of NLRP1B in intestinal organoids.

As *Nlrp1b* is polymorphic, we generated organoids from WT BALB/cJ mice to determine if allele 1 had greater functionality in the epithelium. It has been previously reported that LT triggers NLRP1B-dependent cell death in RAW 264.7 macrophages and BALB/cJ BMDMs ^57^, but LT has never been tested on organoids. While we found that LT was able to enter the cytosol of IECs, as seen by the disappearance of cytosolic MEK1, we did not observe inflammasome activation as measured by GSDMD cleavage and did not detect any NLRP1B in the organoids.

A majority of published work has focused on NLRP1B in macrophages, which inhabit a relatively sterile environment, and readily undergo pyroptosis in response to bacterial stimulus. In contrast, IECs are frequently exposed to microbes, but are more refractory to innate immune signalling, as evidenced by the limited array of Toll-like receptors present in the intestinal epithelium, which is likely an evolved trait to prevent barrier dysfunction ^58^. Indeed, IECs appear to contradict several assumptions surrounding inflammasome activation and pyroptotic cell death. For example, the intestines are the dominant source of circulating IL-18, which is produced at baseline and in the absence of overt pyroptosis in healthy intestinal tissue ^59–61^. We therefore explored the idea that VbP was able to permeate the cell membrane of IECs and activate NLRP1B, but that the downstream consequence of activation were not canonical inflammasome formation involving Caspase-1 and GSDMD. While admittedly in contrast to what has typically been reported in the NLRP1B field, we hypothesized that in the unique cellular context of the intestine, other pyroptotic effectors such as GSDME or GSDMC could be activated by NLRP1B. GSDME has been shown to elicit pyroptosis in IECs ^62^ and GSDMC is similarly induced by enteric parasites and type 2 cytokines ^39, 40^, making both rational substrates of NLRP1B inflammasome activation. However, we did not observe GSDME (p35) or GSDMC (p30) activation with VbP stimulation in IECs, or activation of respective, upstream caspases.

Since WT organoids from C57BL/6J mice express *Nlrp1b* and yet did not respond to VbP, there remains a possibility that the NLRP1B activation requires specific activating signals. Therefore, we attempted to prime SI organoids with cytokines that recapitulate an immune response and would be released by immune cells of the lamina propria, and found that IL-13, which is heavily involved in the immune response against enteric parasites, upregulated *Nlrp1b* expression. However, IL-13 did not sensitize NLRP1B to VbP, but it is possible that uncharacterized other stimuli might be required.

Notably, *Nlrp1b^-/-^* mice have impaired intestinal T-cell responses during protozoan *Tritrichomonas* infection ^17^ and blunted IL-1β processing in response to *Toxoplasma gondii,* suggesting that NLRP1B is functional in the gut and may regulate anti-parasitic responses ^15, 63^. Compellingly, we found that NLRP1B protein and *Nlrp1b* transcript were induced by *Tritrichomonas* infection in the small intestine. In concert with our in vitro IL-13 data, these results point towards a yet unknown epithelial intrinsic role for this NLR during parasite infection.

While we found that NLRP1B was expressed in C57BL/6J intestinal organoids, it remains possible that the defective response to VbP in this tissue could be, at least in part, caused by defective expression or function of the close paralog NLRP1A, as compared to BMDMs. Indeed, previous work has demonstrated that NLRP1A can also respond to VbP ^13^. Notably, while *Nlrp1b* is expressed in both BMDMs and ileal organoids from C57BL/6J animals, *Nlrp1a* was absent from the epithelial compartment. It is therefore plausible that *Nlrp1a* expression is required to sensitive cells to VbP, where absence of NLRP1A in the intestinal epithelium renders the tissue refractory to stimulus, despite the presence of NLRP1B. Importantly, evidence supporting a role for NLRP1A in the response to VbP is primarily derived from experiments performed in recombinant systems, using over-expressed constructs of NLRP1 alleles or with mouse strains which are naturally deficient in either NLRP1A or NLRP1B alleles ^13^. As such, the relative contribution of endogenous NLRP1A, when present in tissue expressing functional NLRP1B, as is the case in C57BL/6J BMDMs, has not been determined. Indeed, it is currently unclear to what extent the response to VbP in C57BL/6J-derived BMDMs is driven by either NLRP1B or the NLRP1A paralog, or by a combination of both, since the *Nlrp1b^-/-^* mice used in this study (which are commercially available and commonly used for NLRP1B-associated studies in mice) also harbour deletion of *Nlrp1a*, which is immediately adjacent to the *Nlrp1b* locus. Likewise, while *Nlrp1a* is absent in C57BL/6J ileal organoids at baseline, there remains the possibility that it can be induced with the correct stimulus. To what extent transcriptional induction of NLRP1 alleles correlates to functional PRR responses is also unknown. Conversely, whether the reduced expression of *Nlrp1a* in the epithelium represents a tissue-specific adaptation, required for normal homeostasis remains to be determined. Further studies, entailing the generation of C57BL/6J mice specifically deficient for either *Nlrp1a* or *Nlrp1b* will be required to address the relative contribution of each paralog in the response to VbP.

In sum, while it is surprising that we were unable to activate NLRP1B in organoids, our data suggest a model wherein tissue specific adaptations to PRR responses may have placed epithelial NLRP1B under yet unknown regulatory mechanisms in order to restrain inflammasome activity in the intestine. Further research is required to delineate these mechanisms and more fully characterize the role of NLRP1B in the epithelium.

## ACKNOWLEDGEMENTS

We thank Dr. Mohanish Deshmukh (University of North Carolina, Chapel Hill) for providing the *Nlrp1b^-/-^*mice. This work was supported by Canadian Institutes for Health Research (CIHR) and the Crohn’s Colitis Canada (CCC) grants to SEG. NJW was supported by a Canadian Graduate Scholarship (CGS-D).

